# Ca^2+^ and cAMP open differentially dilating synaptic fusion pores

**DOI:** 10.1101/2022.12.24.521856

**Authors:** Dinara Bulgari, Samantha L. Cavolo, Brigitte F. Schmidt, Katherine Buchan, Marcel P. Bruchez, David L. Deitcher, Edwin S. Levitan

## Abstract

Neuronal dense-core vesicles (DCVs) contain neuropeptides and much larger proteins that affect synaptic growth and plasticity. Rather than using full collapse exocytosis that is common in endocrine cells, DCVs at a native intact synapse, the *Drosophila* neuromuscular junction, release their contents via fusion pores formed by kiss and run exocytosis. Here fluorogen activating protein (FAP) imaging reveals the permeability range of synaptic DCV fusion pores and then shows that this constraint is circumvented by cAMP-induced extra fusions with dilating pores that result in DCV emptying. These Ca^2+^-independent full fusions require PKA-R2, a PKA phosphorylation site on the fusion clamp protein complexin and the acute presynaptic function of Rugose/Neurobeachin, a PKA-R2 anchor implicated in learning and autism. Therefore, localized Ca^2+^-independent cAMP signaling opens dilating fusion pores to release large cargo proteins that cannot pass through the narrower fusion pores that normally dominate spontaneous and Ca^2+^-evoked synaptic protein release. Hence, two independent exocytosis triggers (Ca^2+^ and cAMP) vary the composition of released proteins at the synapse by differentially adjusting DCV fusion pores.

## INTRODUCTION

Neuronal dense-core vesicles (DCVs) contain a wide variety of secretory protein cargos including neuropeptides (with molecular weights of 0.6-30 kDa in *Drosophila*) and proteases that function in synaptic plasticity and growth (e.g., 70 kDa tissue plasminogen activator) (Huang *et al*., 1996; Baranes *et al*., 1998). Early studies of secretion from endocrine cells emphasized Ca^2+^-induced DCV emptying by full collapse exocytosis that follows fusion pore dilation. However, studies of spontaneous and activity-evoked release by DCVs in the intact *Drosophila* neuromuscular junction (NMJ) with two different imaging approaches failed to detect DCV emptying and instead produced data consistent with release via fusion pores formed by kiss and run exocytosis (Wong *et al*., 2015; Bulgari *et al*., 2019). Partial release by kiss and run exocytosis of presynaptic DCVs is conducive with the cell biology of neurons: because these organelles are replaced by axonal transport that can take days, kiss and run exocytosis may prevent rapid depletion of stores at distal sites of release that are slow to refill. However, it is not known whether presynaptic fusion pores allow for release of large DCV cargos.

Imaging permeation through DCV fusion pores is difficult in native synapses that contain DCVs and small synaptic vesicles (SSVs) because fluid phase fluorophores used with endocrine cells (e.g., Takahashi *et al*., 2002) cannot distinguish between the two vesicle types and cutting-edge superresolution microscopy methods that resolve fusion pore dilation rely on *in vitro* preparations (e.g., Anantharam *et al*., 2011). Likewise, electrical and electrochemical fusion pore measurements (Sharma and Lindau, 2018) are not well suited to typical native cotransmitting synaptic boutons. However, synaptic DCV fusion pores were detected recently using membrane-impermeant malachite green (MG)-based fluorogens and a fluorogen activating protein (FAP) targeted to the DCV lumen so that, upon opening of a fusion pore, fluorogens pass through the fusion pore into the DCV to bind the FAP with picomolar affinity and generate far-red fluorescence (Bulgari *et al*., 2019). Because previously used fluorogens varied in shape and constituent moieties, and their hydrodynamic sizes were not characterized (Bulgari *et al*., 2019), the permeability of presynaptic fusion pores was not studied systematically. Nevertheless, an experimental approach for studying synaptic fusion pore permeability was established.

Here a series of single chain PEGylated membrane impermeant MG fluorogens and a FAP inserted into the neuropeptide *Drosophila* insulin-like peptide 2 (i.e., Dilp2-FAP, Bulgari *et al*., 2019) are used at the *Drosophila* NMJ to show that synaptic fusion pores that open spontaneously and in response to activity are not permeable to large DCV cargos. Then the conundrum of how such proteins can be efficiently released from presynaptic DCVs is resolved by establishing that anchored, activated PKA acts via complexin phosphorylation to open dilating fusion pores that results in DCV emptying.

## RESULTS

### Synaptic DCV fusion pore permeability

Fluorogens were synthesized via conjugating single chain polyethylene glycol (PEG) adducts to MG. Their hydrodynamic sizes were then determined in comparison to proteins by size exclusion fast protein liquid chromatography (FPLC) (see **Methods**). The hydrodynamic properties of these fluorogens scaled with molecular weight differently than proteins; for example, the MG conjugate to a 1 kDa PEG chain with a molecular weight of 1.55 kDa behaved in solution like a 4.3 kDa protein. Because the hydrodynamic size of fluorogens was not taken into account previously, here sizes are expressed in terms of apparent protein molecular weights. Specifically, a series of unbranched MG-PEGs with apparent protein molecular weights ranging from 4.3 to 71.1 kDa were examined for activity-evoked FAP responses at *Drosophila* muscle 6/7 NMJ type Ib boutons. As demonstrated with other membrane impermeant fluorogens such as MG-BTau, FAP responses reflect the opening of fusion pores and the retention of fluorescent MG-Dilp2-FAP complexes that do not exit through fusion pores (Bulgari *et al*., 2019).

Responses to 1 minute of 70 Hz stimulation led to robust labeling with the 4.3 kDa fluorogen, but were attenuated with larger fluorogens (Fig. 1A-D). For the three smaller fluorogens (4.3, 12.5 And 32.4 kDa), the incremental changes could reflect the slower diffusion of larger fluorogens and the effect of approaching the size cutoff of the fusion pore, which is well suited for release of *Drosophila* neuropeptides. However, the two largest fluorogens (53.2 and 71.1 kDa), with incrementally modest increases in hydrodynamic size (because radius is proportional to the cube root of protein molecular weight), barely produced any signals (Fig. 1C,D), thereby indicating poor permeation through synaptic DCV fusion pores that open in response to Ca^2+^ elevation induced by activity. With spontaneous fusion pore openings (Bulgari *et al*., 2019), the same properties are evident: in contrast with the 4.3 and 32.4 kDa fluorogens, time-dependent labeling was not seen with incubation of the NMJ with the 53.2 kDa fluorogen in the absence of Ca^2+^ for 8 minutes (Fig. 1E,F,G). Thus, synaptic fusion pore permeability is sufficient for the release of *Drosophila* neuropeptides, but excludes DCV cargos that exceed the fusion pore cutoff, which falls between 32.4 and 53.2 kDa.

**Fig. 1.**
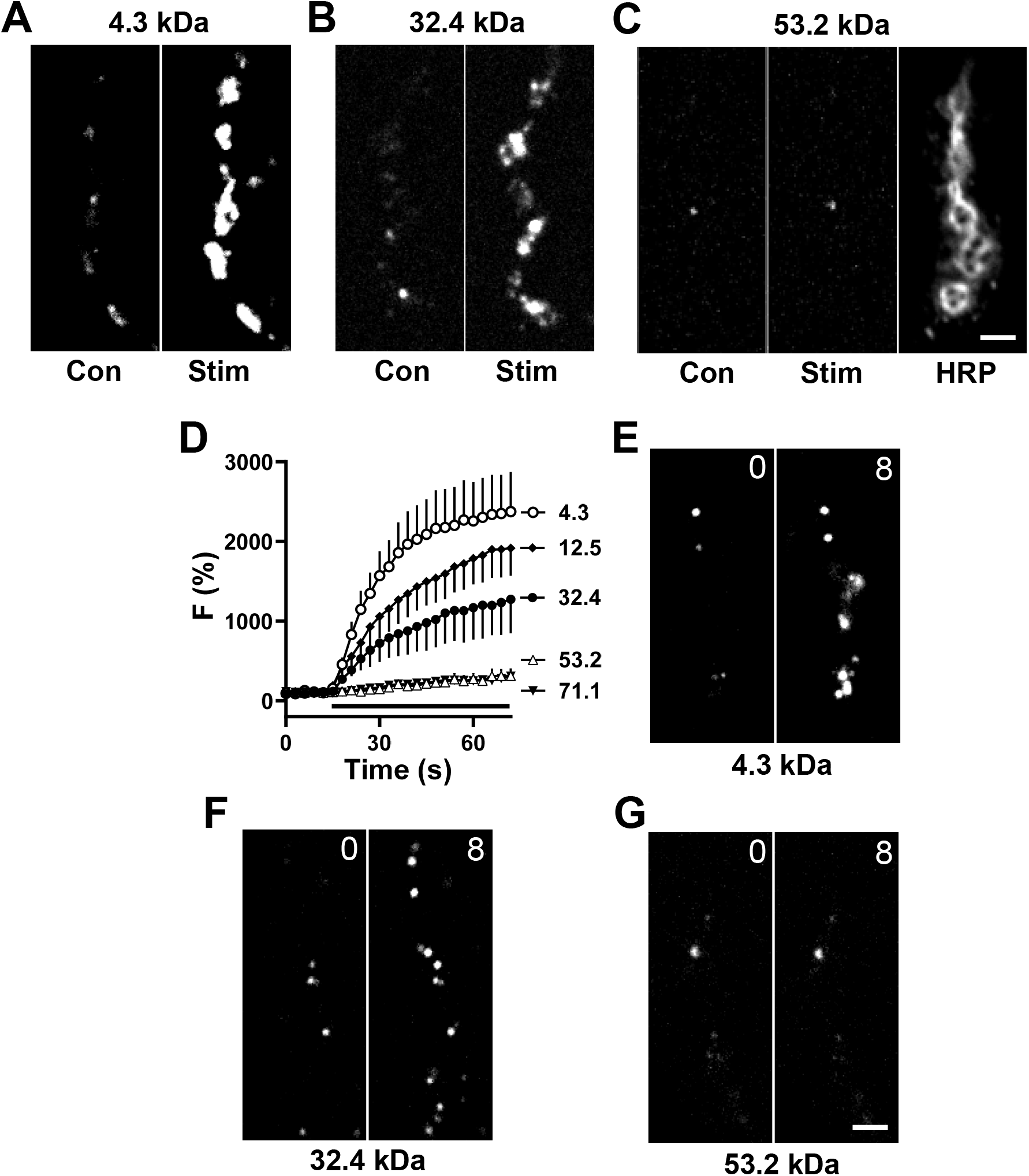
Permeability of PEG derivatives through DCV fusion pores. Contrast-enhanced fluorescent images of type Ib boutons from Ok6-GAL4 UAS-Dilp2FAP larvae before and after 70 Hz stimulation for 60 s in the presence of 4.3 kDa (**A**), 32.4 kDa (**B**) and 53.2 kDa (**C**) MG-PEG dyes, each at 1 µM. Anti-horseradish peroxidase immunofluorescence (HRP) is shown to indicate location of boutons. (**D**) Time-course of activity-evoked FAP responses in the presence of MG-PEG dyes normalized to initial labeling. Apparent protein sizes shown in symbol labels. Black bar, 70 Hz stimulation. 4.3 kDa (n=8), 12.5 kDa (n=7), 32.4 kDa (n=5), 53.2 kDa (n=6), 71.1 kDa (n=6). Contrast-enhanced fluorescent images of Dilp2-FAP expressing boutons in the absence of Ca^2+^ after application of 4.3 kDa (**E**), 32.4 (**F**) and 53.2 kDa (**G**) MG-PEGs. Numbers on the images indicate time in minutes. White scale bars, 2 µm.

### Frequency of DCV emptying events

Given the above synaptic DCV fusion pore permeability, we explored whether NMJ DCVs ever empty to efficiently release large cargos, a process often referred to as full fusion (FF). For this purpose, 1 µM MG-BTau was used to detect spontaneous events that occur in the absence of extracellular Ca^2+^ (Bulgari *et al*., 2019), thus enabling imaging of individual release sites without interference from surrounding events that are stimulated by activity. Single labeling events via kiss and run fusion pores (Fig. 2A, K&R) were readily detected by their sustained labeling and occurred with variable kinetics (Fig. 2B, top and middle); overall, they grew over 23.7 ± 2.7 seconds (n = 37) and could persist for the duration of imaging (up to 4 minutes). However, on rare occasions labeling via the opening of a fusion pore was followed within seconds by the rapid loss of fluorescence (Fig. 2A, FF). Again, time courses were variable (Fig. 2C); the lifetime of these events was 17.25 ± 3.81 seconds with a rise time to peak fluorescence of 5.62 ± 0.53 seconds (n=8). Based on the imaging field of view and the depth of field, such events could not be attributed to undocking and transport of DCVs. Rather, following MG fluorogen influx through the fusion pore and binding to the FAP, MG-FAP complexes that normally are retained (i.e., cannot exit through fusion pores) must have been released by fusion pore dilation that resulted in DCV emptying. Interestingly, such full fusions could occur with a DCV that had already had a kiss and run event (Fig. 2C, bottom). Overall, DCV full fusions only occurred in 3.6% of exocytotic events at the synapse in these experiments (Fig. 3A, Con; filled bar shows kiss and run, open bar shows full fusion; frequencies expressed per bouton).

**Fig. 2.**
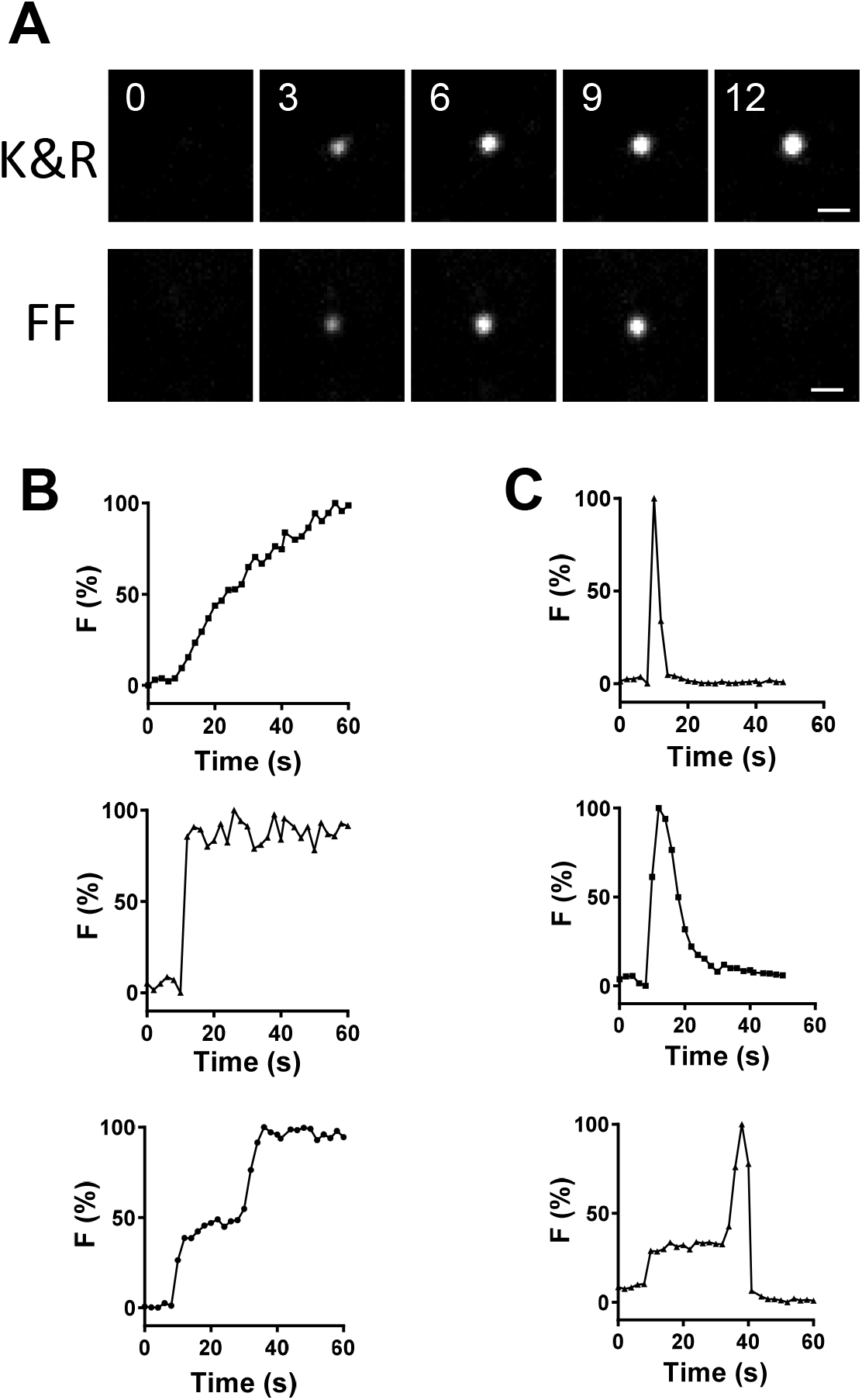
Individual spontaneous kiss and run and full fusion events. (**A**) Contrast-enhanced fluorescent images of single labeled Dilp2-FAP presynaptic sites in the absence of Ca^2+^ after application of MG-BTau fluorogen in OK6>Dilp2-FAP type Ib boutons. Upper panels labeled K&R show consecutive images of kiss and run pore opening, while the lower panels labeled FF show consecutive images of full fusion pore dynamics. Numbers on images indicate times in seconds. Scale bar, 1 µm. Normalized time courses of representative individual (**B**) kiss and run and (**C**) full fusion sites.

**Fig. 3.**
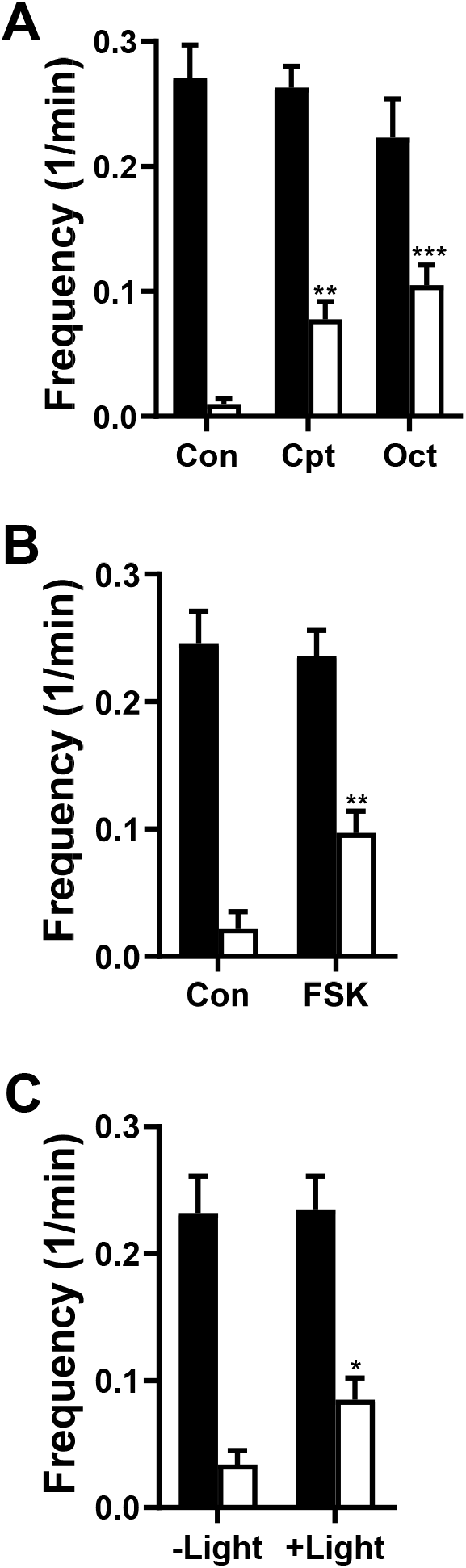
cAMP increases the frequency of spontaneous full fusions. (**A**) Frequency of kiss and run (filled bars) and full fusions (open bars) in control boutons (Con, n=8), boutons treated with the cAMP agonist 8-parachlorophenyl-thio-cAMP (Cpt, n=7) or boutons treated with octopamine (Oct, n=11) in the presence of MG-BTau. Statistical comparisons are always between like colored bars; **p < 0.01, ***p < 0.001, Dunnett’s posttest following one-way ANOVA. (**B**) Frequency of kiss and run (filled bars) and full fusions (open bars) per bouton in DMSO control (Con, n=12) and FSK treated neurons (FSK, n=17) in the presence of MG-BTau, **p < 0.01, unpaired t-test. (**C**) Frequency of kiss and run (filled bars) and full fusions (open bars) in PACα expressing boutons without illumination (-Light, n=11) and light-activated PACα (+Light, n=16) in the presence of MG-BTau, Statistical comparisons are always between like colored bars; *p < 0.05, unpaired t-test. There were no significant differences between kiss and run fusions frequency (filled bars) in presented experiments, unpaired t-test. All data derived from Ib boutons of OK6>Dilp2-FAP animals.

### cAMP evokes opening of dilating fusion pores

Because it seems unlikely that such rare full fusions wholly account for release of large DCV cargo proteins, we considered that signaling might trigger synaptic DCV full fusions. A candidate for such a trigger is cAMP, which evokes Ca^2+^ independent release from DCVs at the *Drosophila* NMJ (Shakiryanova *et al*., 2011; Bulgari *et al*., 2018). Therefore, we tested whether cAMP in the absence of extracellular Ca^2+^ affects the frequency of kiss and run and/or full fusions.

Four independent experimental approaches showed that cAMP selectively evokes presynaptic DCV full fusions. First, the membrane permeant cAMP analog 8-parachlorophenyl-thio-cAMP (CPT, 1 mM) increased the frequency of full fusions without affecting kiss and run events (Fig. 3A). Second, bath application of 100 µM octopamine, an insect neuromodulator that acts via cAMP to promote the growth of NMJ arbors (Koon *et al*., 2011), also increased the frequency of DCV emptying with no effect on the kiss and run event frequency (Fig. 3A, Oct). Third, bath application of 100 µM forskolin (FSK), an adenylate cyclase activator, increased the full fusion frequency without altering kiss and run fusions frequency (Fig. 3B) or the rise times of responses (25.5 ± 3 s (n = 23) for kiss and run; 6.0 ± 1.0 (n = 11) for full fusions). Thus, frequency of one subtype of release is affected without changing kinetics of individual events. Finally, following motor neuron specific expression of the photoactivatable adenylate cyclase PACα (Schröder-Lang et al, 2007), blue light activation of the adenylate cyclase increased the frequency of DCV full fusions, again with no change in kiss and run frequency (Fig. 3C). Together, these results reveal that cAMP evokes release from presynaptic DCVs by selectively increasing the frequency of full fusions without affecting ongoing kiss and run exocytosis.

### Anchored PKA-R2 is required for cAMP-evoked full fusions

The effects of cAMP are mediated by two ubiquitously expressed intracellular cAMP effectors, protein kinase A (PKA) and the exchange protein directly activated by cAMP (Epac) (de Rooij, 1998). To test for a role of Epac, we began with bath application of the Epac activator 8-(4-methoxyphenylthio)-2′-O-methyl-cAMP (Me, 200 µM). However, no changes in the frequencies of emptying or kiss and run events were produced (Fig. 4A,D). Furthermore, pretreatment of boutons with Epac inhibitor ESI-09 (10 µM) did not disrupt the forskolin-evoked increase in full fusion frequency (Fig. 4B) or alter kiss and run events (Fig. 4E). Finally, in Epac null animals (Shakiryanova *et al*., 2011), which have a P element insertion near the C-terminal cyclic nucleotide-binding domain, the selective effect of forskolin on emptying persisted (Fig. 4C,F). Interestingly, the latter two approaches to reducing epac function tended to increase basal emptying frequency without eliminating the forskolin effect. This suggests that epac may constrain fusion pore dilation in the synaptic terminal independently of cAMP. But more important here, Epac is not required for cAMP-induced DCV full fusions.

**Fig. 4.**
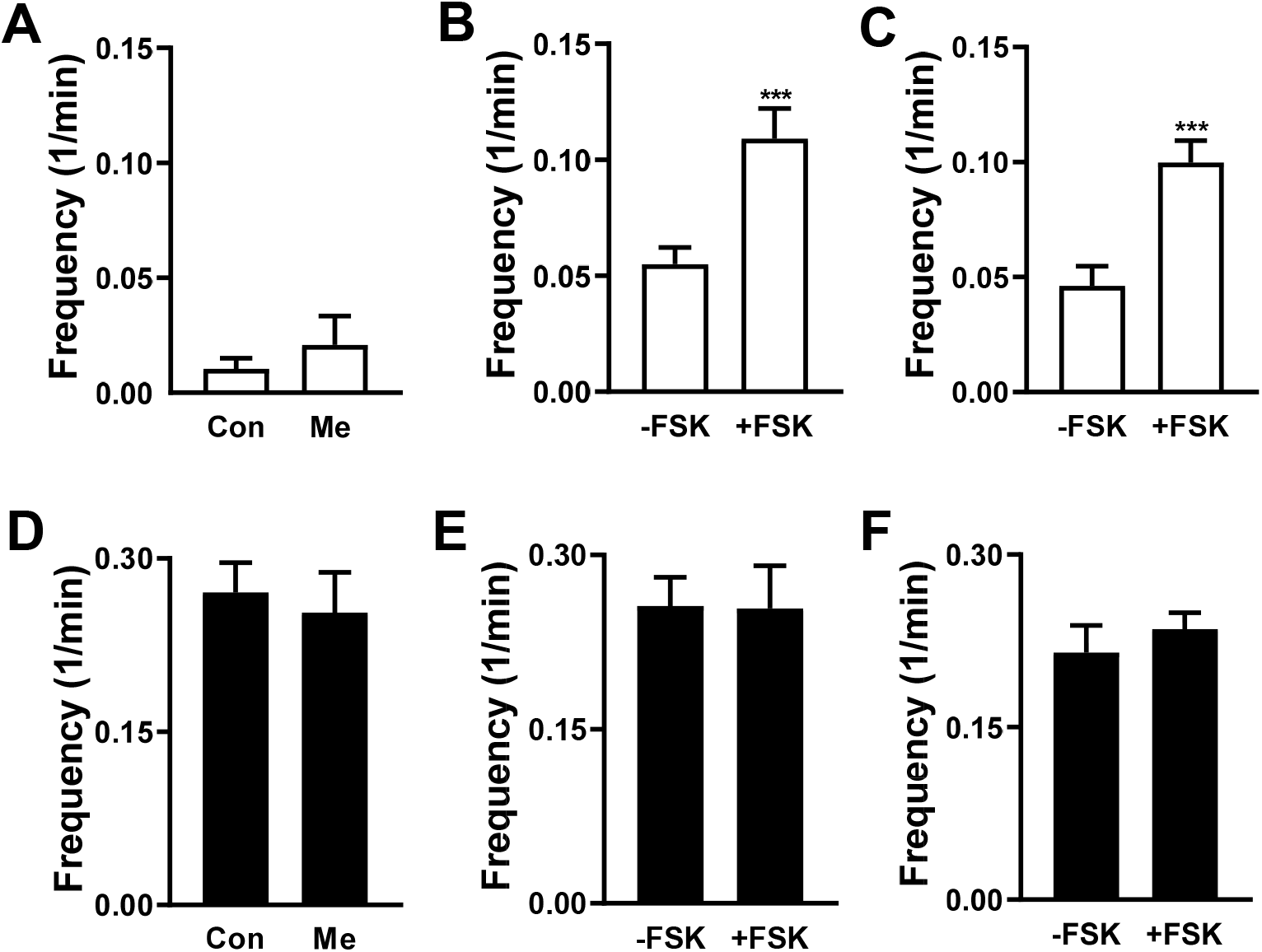
Epac is not involved in cAMP-induced full fusions. (**A**) Frequency of full fusions in presence of MG-BTau in control OK6>Dilp2-FAP boutons (Con, n=12) and pretreated with Epac activator 8-(4-methoxyphenylthio)-2′-O-methyl-cAMP (Me, n=7). No significant difference was detected with unpaired t-test. (**B**) Epac inhibitor ESI-09 without FSK (-FSK, n=10,) and with FSK (+FSK, n=7), ***p < 0.001. (**C**) in flies expressing Epac null without FSK (-FSK, n=11) and with FSK (+FSK, n=17), ***p < 0.001. (**D-F**) Frequency of kiss and run fusions with conditions corresponding to **A-C**. Note that there were no significant differences in kiss and run fusions for the experimental conditions presented here.

In contrast, presynaptic PKA containing the type 2 regulatory subunit (PKA-R2) is required for cAMP-evoked DCV full fusions. First, the PKA catalytic subunit inhibitor H89 (20 µM) blocks the forskolin-induced full fusions (Fig. 5A, open bars) without altering kiss and run events (Fig. 5A, filled bars). Second, presynaptic expression of a PKA-R1 dominant negative subunit failed to affect the cAMP effect (Fig. 5B), consistent with the involvement of PKA-R2. Third, knockdown of PKA-R2 inhibited the cAMP effect. For these experiments, initial attempts to express Dilp2-FAP and a PKA-R2 RNAi with the motor neuron driver OK6-GAL4 did not produce viable progeny. Therefore, PKA-R2 RNAi was expressed only in type Ib boutons with the Dip-β-GAL4 driver (Wang *et al*, 2022). Strikingly, the forskolin effect on emptying frequency was eliminated, while kiss and run events were unchanged (Fig. 5C,D). We also tested the role of PKA-R2 in type Is boutons using the ShakB-GAL4 driver. Forskolin produced the same phenomenon of increased emptying frequency in type Is boutons as found in Ib boutons (Fig. 5E) and this effect was also inhibited by PKA-R2 RNAi knockdown, again without affecting kiss and run events (Fig. 5F). Finally, we focused on the association of PKA-R2 with AKAPs (A-kinase anchoring proteins). Acute treatment of NMJs with 20 µM St-Ht31, a stearated membrane permeant inhibitor of the interaction of PKA-R2 with AKAPs (Vijayaraghavan *et al*., 1997), abolished the facilitating effect of FSK on DCV full fusions with no effect on kiss and run frequency (Fig. 5G). This short-term experiment further implicates PKA-R2 without confounding developmental effects. However, this result cannot localize the interaction of PKA-R2 and AKAP interaction to the presynaptic neuron (as opposed to the postsynaptic muscle). Therefore, RNAi knockdown of *Drosophila* Rugose (also called Akap550), a neuronal PKA-R2 anchor (Han *et al*., 1997), was induced in motor neurons. This genetic presynaptic perturbation inhibited the effect of cAMP on DCV full fusions, again without affecting kiss and run exocytosis frequency (Fig. 5H). Taken together, these experimental results demonstrate that cAMP activation of anchored PKA-R2 triggers Ca^2+^-independent full fusion of presynaptic DCVs.

**Fig. 5.**
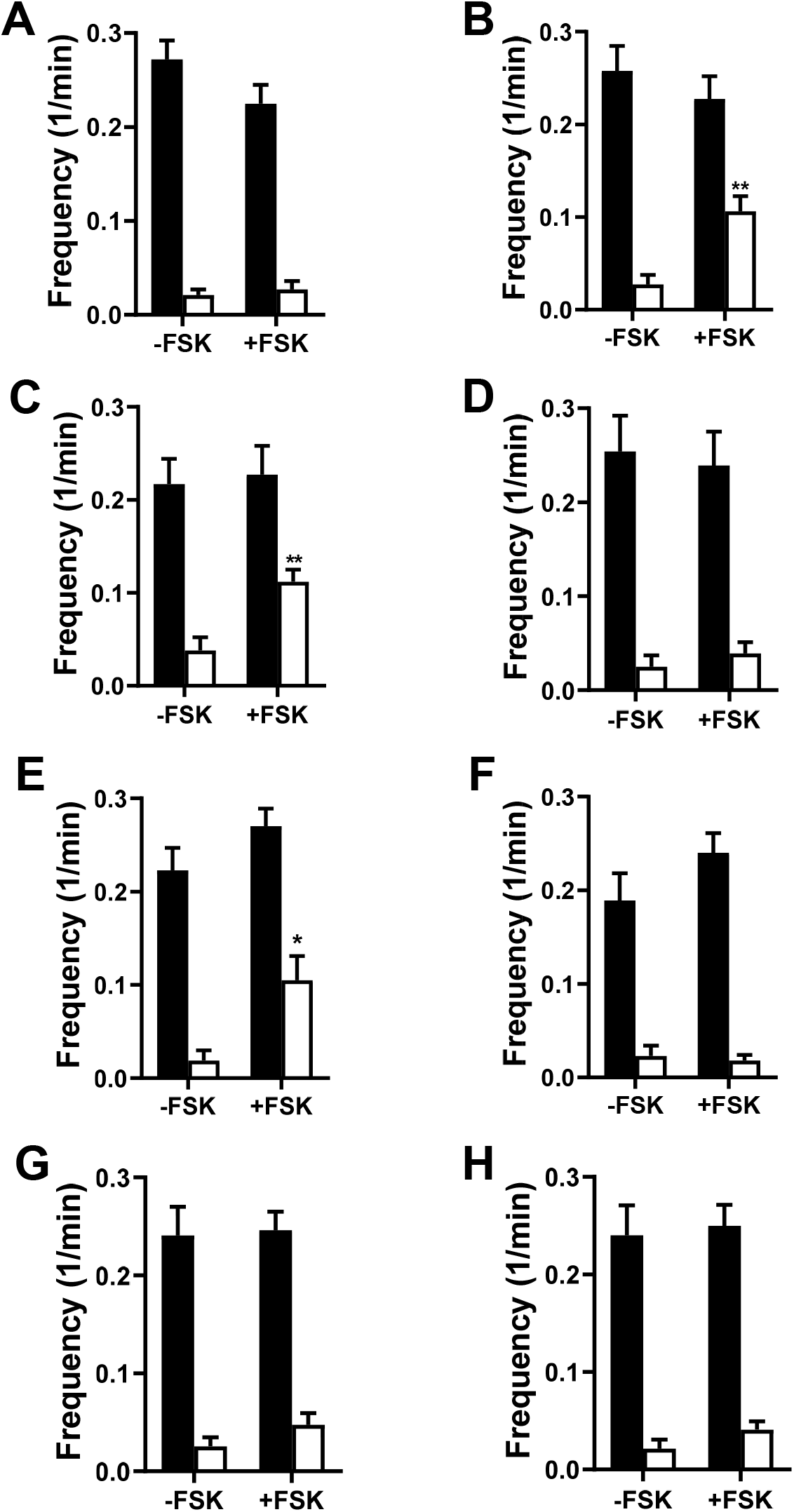
cAMP-induced full fusions require anchored PKA-R2. (**A**) Frequency of kiss and run (filled columns) and full fusions (open columns) in OK6>Dilp2-FAP boutons treated with H89 without (-FSK, n=14) and with FSK application (+FSK, n=9). (**B**) Frequency of fusions in OK6>Dilp2-FAP boutons coexpressing dominant-negative PKA-R1 (-FSK, n=8; +FSK, n=14) **p < 0.01. Frequency of fusions in (**C**) DIPβ-GAL4>Dilp2-FAP boutons (n=8; +FSK, n=9, **p < 0.01) and (**D**) DIPβ-GAL4>Dilp2-FAP boutons also expressing PKA-R2 RNAi (-FSK, n=7; +FSK, n=7). (**E**) ShakB-GAL4>Dilp2-FAP coexpressing the UAS-Valium20 RNAi control with and without FSK treatment (-FSK, n=8; +FSK, n=10) and (**F**) ShakB-GAL4>Dilp2-FAP boutons coexpressing UAS-PKA-R2 RNAi with and without FSK treatment (-FSK, n=8; +FSK, n=19). (**G**) Frequency of fusions in St-Ht31 pretreated OK6>Dilp2-FAP boutons with and without FSK treatment (-FSK, n=12), (+FSK, n=20). (**H**) Frequency of fusions in OK6>Dilp2-FAP boutons coexpressing UAS-rugose with and without FSK (-FSK, n=13; +FSK, n=11). No significant difference was found between kiss and run fusions frequency in FSK-treated and non-treated boutons, unpaired t-test.

### Complexin participates in cAMP-evoked release from synaptic DCVs

The above results pose the question of how rugose-anchored PKA activity induces Ca^2+^-independent synaptic DCV full fusions. Recent findings directed our attention to complexin (Cpx), which is often referred to as fusion clamp because it reduces spontaneous SSV fusion. First, spontaneous release from DCVs and SSVs are similar at the NMJ: both persist in the presence of tetanus toxin and share SNARE dependence (Bulgari *et al*., 2019). Furthermore, *Drosophila* complexin is phosphorylated by PKA at S126 to upregulate (or unclamp) spontaneous SSV release events (Cho *et al*., 2015). Therefore, the increase in spontaneous DCV release events with cAMP could be accounted for by complexin phosphorylation. Second, *in vitro* reconstitution experiments devoid of much of the cellular exocytosis machinery suggest the potential for facilitation of fusion pore dilation by complexin (Pierson and Shin, 2021). Together, these observations support the hypothesis that complexin S126 is required for Ca^2+^-independent PKA-induced spontaneous DCV full fusions.

Therefore, we examined the effect of replacing native complexin with the unphosphorylatable S126A mutant (Cpx^S126A^) by rescuing a Cpx null mutant with neuronal expression of Cpx^S126A^ (i.e., as in Cho *et al*., 2015). First, to test whether the mutant rescue was effective for DCVs, a GFP-tagged neuropeptide (ANF-GFP) (Rao *et al*., 2001) that reports native DCV-mediated release (e.g., Husain and Ewer, 2004) was expressed while replacing Cpx with Cpx^S126A^ and release was measured as the loss of ANF-GFP fluorescence. In the presence of extracellular Ca^2+^, activity-evoked DCV-mediated release in Cpx^S126A^ animals was intact: the ∼20% release (Fig. 6A), followed by some recovery due to activity-dependent capture, matches prior control responses with GFP imaging (Shakiryanova *et al*., 2005, 2006; Wong *et al*., 2015; Cavolo *et al*., 2016). Thus, the Cpx^S126A^ complexin mutant supports Ca^2+^ dependent synaptic DCV exocytosis.

**Fig. 6.**
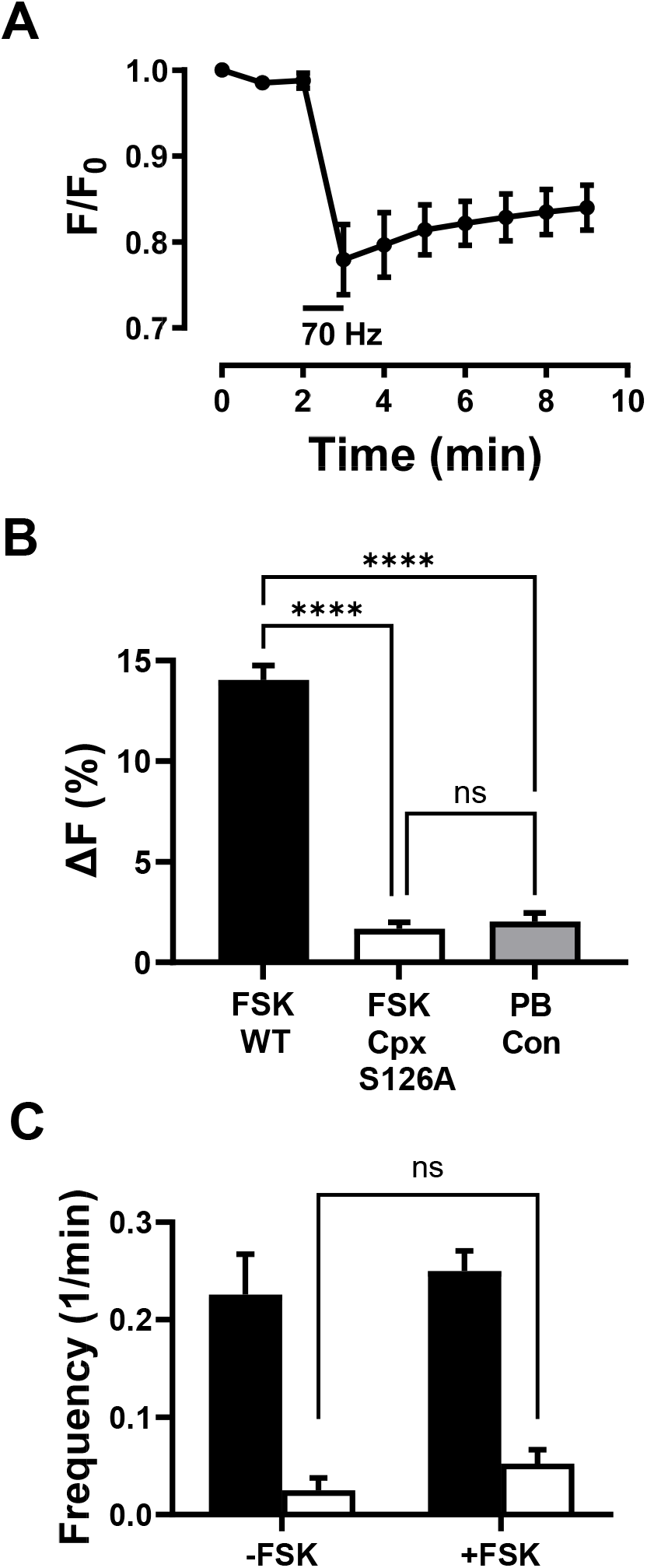
The complexin S126 PKA phosphorylation site is required for cAMP-evoked full fusions and neuropeptide release. Release indicated by loss of ANF-GFP fluorescence is induced by a 1 minute 70 Hz stimulation (indicated by the horizontal bar) in elav-GAL4 UAS-ANF-GFP;;UAS-Cpx^S126A^, Cpx^SH1^ animals (n=4). (**B**) Forskolin evoked ANF-GFP release in 7 minutes is abolished in elav-GAL4 UAS-ANF-GFP;;UAS-Cpx^S126A^, Cpx^SH1^ animals. WT, wild type Cpx; PB Con, photobleaching control. n=4-6, ****p < 0.0001, Tukey’s test following ANOVA. (**C**) Frequency of kiss and run (filled columns) and full fusions (open columns) in boutons from elav-GAL4;UAS-Dilp2-FAP;UAS-Cpx^S126A^, Cpx^SH1^/Cpx^Delta2^ flies in the absence and presence of forskolin (-FSK (n=7) and +FSK (n=14), respectively). Cpx^SH1^ and Cpx^Delta2^ are null mutants. Note that forskolin does not have a statistically significant effect (ns, not significant by t-test) on full fusion frequency (open bars). Likewise, no effect was produced on kiss and run frequency.

Then cAMP-evoked release was measured in the absence of extracellular Ca^2+^. Figure 6B shows that the ANF-GFP release response to forskolin is abolished in Cpx^S126A^ animals: with the mutant the minor change in fluorescence that occurred with forskolin matched the photobleaching control. Finally, FAP imaging was used to measure the impact of Cpx^S126A^ on DCV full fusions (Fig. 6C). As expected from all previous experiments, no change occurred in kiss and run frequency. Furthermore, in accordance with ANF-GFP experiments, cAMP stimulation of DCV full fusions was abolished (i.e., the open bars in Fig. 5C were not significantly different, ns). Therefore, reminiscent of SSVs, complexin’s PKA phosphorylation site is needed for the cAMP-induced increase in frequency of spontaneous DCV-mediated release. However, complexin S126 has the added impact for DCVs of supporting cAMP-evoked opening of dilating fusion pores to enable release of proteins that are too large to permeate through the fusion pores that normally dominate release.

## DISCUSSION

Neuronal DCVs are distinguished by their small size relative to the best studied endocrine DCVs, widely varying contents within a single cell and, at a native intact synapse, the nearly exclusive use of fusion pores formed by dynamin dependent kiss and run exocytosis for spontaneous and activity-evoked release (Wong *et al*., 2015; Bulgari *et al*., 2019, Fig. 3A). The resultant partial release from DCVs fits with the need to preserve presynaptic DCVs, which can only be replaced very slowly by axonal transport, but posed the question of whether large DCV cargos can permeate through fusion pores. FAP imaging with a series of PEGylated fluorogens provided a means to determine the conduction properties of synaptic DCV fusion pores without cargo specific confounds that affect release (e.g., binding to luminal DCV constituents). The experimental data presented here establish that presynaptic DCV fusion pores are large enough to release all *Drosophila* neuropeptides, but insufficient for large proteases that undergo activity-dependent synaptic release (e.g., TPA and neurotrypsin). However, this limit is bypassed by cAMP-induced Ca^2+^-independent full fusions. The resultant efficient release of large cargos may contribute to developmental growth and refinement of NMJ terminals, which depend on cAMP (Zhong *et al*., 1992; Koon *et al*., 2011; Vonhoff and Keshishian, 2017).

cAMP-evoked DCV emptying (Fig. 3) requires PKA-R2 and its anchor Rugose (Fig. 5). cAMP also stimulates spontaneous release from SSVs at the *Drosophila* NMJ (Yoshihara *et al*., 1999; Zhang *et al*., 1999; Cho *et al*., 2015), but the role of Rugose has not been examined for SSVs. However, Rugose affects larval NMJ morphology and a variety of adult behaviors including associative learning (Volders *et al*., 2012; Wise *et al*., 2015). Because Rugose and its mammalian ortholog Neurobeachin, a candidate autism gene (Castermans *et al*., 2003), are localized near the trans Golgi network (Wang *et al*., 2000; Volders *et al*., 2012), effects on synapses and behavior have been proposed to originate from perturbing the somatodendritic compartment (e.g., by affecting Golgi function, postsynaptic receptors and dendritic spines) (Niesmann *et al*., 2011; Volders *et al*., 2012; Gromova *et al*., 2018; Repetto *et al*., 2018). However, the effect of acute disruption of the PKA-R2 anchoring on cAMP-induced full fusions at boutons (Fig. 5G) establishes that Rugose is present and functions in presynaptic boutons. Thus, cAMP-induced spontaneous synaptic release by DCVs and possibly SSVs involving Rugose/Neurobeachin-anchored PKA-R2 may locally control synaptic release to influence behavior.

One way the regulation of fusion could be represented is with a channel gating model in which the fusion pore moves between closed (C), open (O) and dilated (D) states (C ⇌O → D). In this scheme, reaching only the O state results in kiss and run release, while reaching the D state produces full fusion (i.e., DCV emptying). If the cAMP/PKA/complexin pathway selectively increased the number of fusions by promoting the C → O step, the frequency of kiss and run events, as well as subsequent emptying events, would have increased. Yet kiss and run frequency did not increase in response to cAMP, thus excluding this possibility. An alternative hypothesis is that cAMP/PKA/complexin pathway increases the incidence of dilation, which initiates full fusion (the O → D step). With this mechanism, would-be kiss and run release events are converted to full fusions. Therefore, the greater frequency of full fusions would be accompanied with a reduced frequency of kiss and run events. But this was not found, thereby showing cAMP-induced full fusions are not produced at the expense of kiss and run events. Instead, rugose-anchored PKA increases total fusion pore openings with the “extra” events being committed to fusion pore dilation. With simple gating models eliminated, the selective increase in spontaneous full fusions reported here is consistent with complexin phosphorylation both unclamping DCVs normally held in reserve (thus recruiting “extra” DCVs to undergo Ca^2+^-independent exocytosis) and dilating resultant fusion pores. With pore dilation, large cargos that do not readily pass through activity-evoked fusion pores are released. Thus, diverse triggers (Ca^2+^ elevation induced by activity and complexin phosphorylation induced by rugose-anchored PKA) evoke release of distinct DCV cargos by opening differentially dilated fusion pores.

## MATERIALS AND METHODS

### Imaging

*Drosophila melanogaster* third instar larvae were filleted and imaged in Ca^2+^-free HL3 in which 1.5 mM Ca^2+^ was substituted with 0.5 mM EGTA, a Ca^2+^ chelator (70 mM NaCl, 5 mM KCl, 0.5 mM Na_3_EGTA, 20 mM MgCl_2_, 10 mM NaHCO_3_, 5 mM trehalose, 115 mM sucrose, 5 mM hemi-sodium HEPES, pH 7.25). For electrical stimulation experiments they were transferred to HL3 saline that contained (in mM) 70 mM NaCl, 5 KCl, 1.5 CaCl_2_, 20 MgCl_2_, 10 NaHCO_3_, 5 trehalose, 115 sucrose, and 5 hemi-sodium HEPES, pH 7.25 supplemented with 10 mM L-glutamate to prevent muscle contractions. Nerve terminals were stimulated at 70 Hz for 1 minute via segmental nerves with a suction electrode (Shakiryanova et al, 2005). Dilp2-FAP data were acquired in muscle 6/7 type I boutons of with an upright Olympus microscope equipped with a 60×1.1 NA water immersion objective, a Yokogawa CSU-X1 spinning disk confocal head, a Coherent Obis 640 nm laser for fluorogen illumination, a Ludl filter wheel with an ET 700/75 emission filter and a Teledyne Photometrics Prime 95B sCMOS camera. ANF-GFP type Ib bouton data were acquired as described previously (Levitan et al, 2007; Wong *et al*., 2015). PACα photoactivation in synaptic boutons was achieved by illuminating synaptic boutons with 488 nm light for 300 ms at 0.33 Hz for the whole duration of FAP imaging (4 minutes). Quantification of fluorescence intensity was performed with ImageJ software (https://imagej.nih.gov/ij/) as previously described (Levitan et al, 2007). Spontaneous fusion events were counted manually from time-lapse images taken for 4 min at 0.33 or 0.5 Hz. Statistical analysis and graphing were performed with GraphPad Prism software. Error bars represent SEM. Statistical significance was determined with Student’s t-test for two experimental groups and ANOVA with Dunnett’s or Tukey’s post-test for more experimental groups.

### Reagents

For synthesis of MG-PEGs, commercially available mPEG-azides of sizes 1K, 2K, 5K, 10K and 20 K (Biochempeg Scientific Inc.) were coupled to MG-EDA-alkyne via click reaction (Szent-Gyorgyi *et al*., 2023). Hydrodynamic sizes in terms of apparent protein molecular weights were determined by FPLC with a HiPrep 26/60 Sephacryl S-200 high-resolution column with protein standards ranging from 6.5 to 66.5 kDa (aprotinin, cytochrome C, carbonic anhydrase, albumin). Resultant apparent protein molecular weights were: 4.3 kDa for MG-PEG1K, 12.5 kDa for MG-PEG2K, 32.4 kDa for MG-PEG5K, 53.2 kDa for MG-PEG10K and 71.1 kDa for MG-PEG20K. MG-PEG conjugates and MG-BTau were used at 1 µM final concentration in imaging experiments. Octopamine and MG-BTau were bath applied immediately before imaging. Time-lapse images were taken for 4 minutes at 0.3 or 0.5 Hz. Boutons were treated with 8-parachlorophenyl-thio-cAMP and 8-(4-methoxyphenylthio)-2′-O-methyl-cAMP for 15 minutes before application of 1 µM MG-BTau followed by image sequence acquisition. stHT-31, ESI-09 and H-89 were applied for 15 to 20 min before application of forskolin, then imaged 3 minutes after forskolin exposure in the presence of MG-BTau. Forskolin was dissolved in DMSO as stock and subsequently diluted to yield a final concentration of DMSO of 0.1%. An equivalent amount of the vehicle was used as the matched control. Forskolin, 8-parachlorophenyl-thio-cAMP and ESI-09 were from Sigma, 8-(4-methoxyphenylthio) -2′-O-methyl-cAMP from Biolog Life Sciences, H-89 from Cayman Chemicals and St-Ht31 from Fisher Scientific.

### Flies

UAS-Dilp2-FAP and UAS-ANF-GFP flies were described previously (Rao *et al*., 2001; Bulgari *et al*., 2019). Epac function was disrupted based a P-element insertion in the gene (Epac KG00434; Bloomington stock #13663). Other Bloomington stocks used were: OK6-GAL4 (#64199), DIPβ-GAL4 (#90316), UAS-PACα (#78790), the dominant negative PKA-R1 UAS-PKA.R1BDK (#35550), UAS-PKA-R2 RNAi (#53930), Valium20 RNAi control (#35783), UAS-rugose RNAi (#57703) and the complexin null Cpx^Delta2^ (#64252). The ShakB-GAL4 line was from Tanja Godenschwege (Florida Atlantic University). J. Troy Littleton (MIT) provided elav-GAL4;;UAS-Cpx^S126A^, Cpx^SH1^/TM6B,Tb flies, with Cpx^SH1^ being a complexin null mutant.

## Acknowledgements

We thank Drs. Tanja Godenschwege (Florida Atlantic University), Robert A. Carrillo (University of Chicago) and J. Troy Littleton (Massachusetts Institute of Technology) for flies, and Drs. Manfred Lindau (University of Miami), Ronald Holz (University of Michigan) and Christopher Szent-Gyorgyi (Carnegie Mellon University) for helpful discussions.

This research was supported by the National Institutes of Health grant R01 NS032385 to ESL.

## REFERENCES

Anantharam A, Bittner MA, Aikman RL, Stuenkel EL, Schmid SL, Axelrod D, Holz RW. A new role for the dynamin GTPase in the regulation of fusion pore expansion. Mol Biol Cell. 22, 1907–18 (2011).

Baranes D, et al. Tissue plasminogen activator contributes to the late phase of LTP and to synaptic growth in the hippocampal mossy fiber pathway. Neuron 21, 813–25 (1998).

Bulgari D, et al., Activity-evoked and spontaneous opening of synaptic fusion pores. Proc. Natl. Acad. Sci. U.S.A. 116, 17039–17044 (2019).

Bulgari D., Jha A., Deitcher D. L., Levitan E.S. Myopic (HD-PTP, PTPN23) selectively regulates synaptic neuropeptide release. Proc Natl Acad Sci U S A. 115, 1617–1622 (2018).

Castermans D, et al. The neurobeachin gene is disrupted by a translocation in a patient with idiopathic autism. J Med Genet. 40, 352–6 (2003).

Cavolo SL, Bulgari D, Deitcher DL, Levitan ES. Activity Induces Fmr1-Sensitive Synaptic Capture of Anterograde Circulating Neuropeptide Vesicles. J Neurosci. 36, 11781–11787 (2016).

Cho RW, et al. Phosphorylation of Complexin by PKA Regulates Activity-Dependent Spontaneous Neurotransmitter Release and Structural Synaptic Plasticity. Neuron 88, 749–61 (2015).

J de Rooij, et al. Epac is a Rap1 guanine-nucleotide-exchange factor directly activated by cyclic AMP. Nature 396,474–7 (1998).

Gromova KV, et al. Neurobeachin and the kinesin KIF21B are critical of endocytic recycling of NMDA receptor and regulate social behavior. Cell Rep. 23, 2705–2717 (2018).

Han JD, Baker NE, Rubin CS. Molecular characterization of a novel A kinase anchor protein from Drosophila melanogaster. J Biol. Chem. 272, 26611–9 (1997).

Huang YY et al. Mice lacking the gene encoding tissue-type plasminogen activator show a selective interference with late-phase long-term potentiation in both Schaffer collateral and mossy fiber pathways. Proc Natl Acad Sci U S A. 93, 8699–704. (1996).

Husain QM, Ewer J. Use of targetable gfp-tagged neuropeptide for visualizing neuropeptide release following execution of a behavior. J. Neurobiol. 59, 181–91 (2004).

Koon AC, et al. Autoregulatory and paracrine control of synaptic and behavioral plasticity by octopaminergic signaling. Nat Neurosci. 14,190–9 (2011).

Levitan ES, Lanni F, Shakiryanova D. In vivo imaging of vesicle motion and release at the Drosophila neuromuscular junction. Nat Protoc. 2, 1117–25 (2007).

Niemann K, et al. Dendritic spine formation and synaptic function require Neurobeachin. Nat Comm. 2, 557 (2011).

Pierson J, Shin YK. Stabilization of the SNARE core by Complexin-1 facilitates fusion pore expansion. Front. Mol. Biosci. 8, 805000 (2021).

Rao S, Lang C, Levitan ES, Deitcher DL. Visualization of neuropeptide expression, transport, and exocytosis in Drosophila melanogaster. J. Neurobiol. 49, 159–172 (2001).

Repetto D, et al. Molecular dissection of Neurobeachin function at excitatory synapses. Front. Synaptic Neurosci. 10, 28 (2018).

Schröder-Lang S, et al. Fast manipulation of cellular cAMP level by light in vivo Nat Methods 4, 39–42 (2007).

Shakiryanova D, Tully A, Hewes RS, Deitcher DL, Levitan ES. Activity-dependent liberation of synaptic neuropeptide vesicles. Nat Neurosci. 8, 173–8 (2005).

Shakiryanova D, Tully A, Levitan ES. Activity-dependent synaptic capture of transiting peptidergic vesicles. Nat. Neurosci. 9, 896–9000 (2006).

Shakiryanova D., Zettel G. M., Gu T., Hewes R. S., Levitan E. S. Synaptic neuropeptide release induced by octopamine without Ca^2+^ entry into the nerve terminal. Proc Natl Acad Sci U S A. 108, 4477–4481 (2011).

Sharma S, Lindau M. The fusion pore, 60 years after the first cartoon. FEBS Lett. 592, 3542–3562 (2018).

Szent-Gyorgyi C., et al. Bottom-Up design: a modular golden gate assembly platform of yeast plasmids for simultaneous secretion and surface display of distinct FAP fusion proteins. ACS Synth Biol. 11, 3681–3698 (2022).

Takahashi N., Kishimoto T., Nemoto T., Kadowaki T., Kasai H. Fusion pore dynamics and insulin granule exocytosis in the pancreatic islet. Science 297, 1349–52 (2002).

Vijayaraghavan S, Goueli SA, Davey MP, Carr DW. Protein kinase A-anchoring inhibitor peptides arrest mammalian sperm motility. J Biol Chem. 272, 4747–52 (1997).

Volders K, et al. Drosophila rugose is a functional homolog of mammalian Neurobeachin and affects synaptic architecture, brain morphology, and associative learning. J Neurosci. 32, 15193–204 (2012).

Vonhoff F, Keshishian H. Cyclic nucleotide signaling is required during synaptic refinement at the Drosophila neuromuscular junction. Dev. Neurobiol. 77, 39–60 (2017).

Wang X, et al. Neurobeachin: A protein kinase A-anchoring, beige/Chediak-higashi protein homolog implicated in neuronal membrane traffic. J Neurosci. 20, 8551–65 (2000).

Wang Y, et al. Systematic expression profiling of Dpr and DIP genes reveals cell surface codes in Drosophila larval motor and sensory neurons. Development 49, dev200355 (2022).

Wise A, et al. Drosophila mutants of the autism candidate gene neurobeachin (rugose) exhibit neuro-developmental disorders, aberrant synaptic properties, altered locomotion, and impaired adult social behavior and activity patterns. J Neurogenet. 29, 135–43 (2015).

Wong M. Y., Cavolo S. L., Levitan E. S., Synaptic neuropeptide release by dynamin-dependent partial release from circulating vesicles. Mol. Biol. Cell 26, 2466–2474 (2015).

Yoshihara M, et al. Selective effects of neuronal-synaptobrevin mutations on transmitter release evoked by sustained versus transient Ca^2+^ increases and by cAMP. J Neurosci. 19, 2432–41 (1999).

Zhang D, Kuromi H, Kidokoro Y. Activation of metabotropic glutamate receptors enhances synaptic transmission at the Drosophila neuromuscular junction. Neuropharmacology 38, 645–57 (1999).

Zhong Y, Budnik V, Wu CF. Synaptic plasticity in Drosophila memory and hyperexcitable mutants: role of cAMP cascade. J Neurosci.12, 644–51 (1992).

